# A Vector System Encoding Histone H3 Mutants Facilitates Manipulations of the Neuronal Epigenome

**DOI:** 10.1101/2024.01.18.576334

**Authors:** Sophie Warren, Sen Xiong, Daisy Robles, José-Manuel Baizabal

## Abstract

The differentiation of developmental cell lineages is associated with genome-wide modifications in histone H3 methylation. However, the causal role of histone H3 methylation in transcriptional regulation and cell differentiation has been difficult to test in mammals. The experimental overexpression of histone H3 mutants carrying lysine-to-methionine (K-to-M) substitutions has emerged as an alternative tool for inhibiting the endogenous levels of histone H3 methylation at specific lysine residues. Here, we leverage the use of histone K-to-M mutants by creating Enhanced Episomal Vectors that enable the simultaneous depletion of multiple levels of histone H3 lysine 4 (H3K4) or lysine 9 (H3K9) methylation in projection neurons of the mouse cerebral cortex. Our approach also facilitates the simultaneous depletion of H3K9 and H3K27 trimethylation (H3K9me3 and H3K27me3, respectively) in cortical neurons. In addition, we report a tamoxifen-inducible Cre-FLEX system that allows the activation of mutant histones at specific developmental time points or in the adult cortex, leading to the depletion of specific histone marks. The tools presented here can be implemented in other experimental systems, such as human *in vitro* models, to test the combinatorial role of histone methylations in developmental fate decisions and the maintenance of cell identity.

## INTRODUCTION

Cell diversity in the adult organism arises during embryonic development from the activation of cell-type-specific transcriptional programs. Lysine methylation at the amino-terminal region of histone H3 is a chromatin post-translational modification associated with distinct transcriptional states^1^. The “histone code” model postulates that the combinatorial or sequential activity of histone methylations at multiple lysine residues determines the transcriptional output —and thereby the identity— of a cell^2,3^. Notably, genome-wide alterations in histone methylation are associated with human developmental disorders, cancer, and aging^1^. However, testing the individual and combined roles of histone methylations on developmental fate specification, differentiation, and the maintenance of cell identity, remains a major technical roadblock that demands the generation and optimization of novel experimental tools and approaches^4^.

The analysis of gene regulation by histone H3 methylation is substantially complicated by the presence of mono-, di-, and trimethylated lysine residues^1^. In some cases, distinct levels of histone H3 methylation are enriched at different *cis*-regulatory elements in the genome. For instance, histone H3 lysine 4 monomethylation (H3K4me1) preferentially marks active or poised enhancers, whereas lysine 4 trimethylation (H3K4me3) is mostly present at the promoter of actively transcribed genes^5,6^. Nonetheless, it remains unclear whether H3K4 methylation plays a primary role in gene activation^4,7^. In other instances, multiple levels of methylation on a lysine residue might have redundant roles in controlling specific transcriptional outputs. In this regard, histone H3 lysine 9 di- and trimethylation (H3K9me2 and H3K9me3, respectively) are both linked to transcriptional repression^8^. However, the unique and cooperative roles of H3K9me2 and H3K9me3 in gene silencing and their functional crosstalk with other repressive marks, such as histone H3 lysine 27 trimethylation (H3K27me3), remain mostly unknown^9^.

The main experimental approach to study the role of histone methylation in mammalian cells is the generation of loss-of-function mutations in the genes encoding for histone methyltransferases^4^. The main caveat of this approach is that histone methylations are usually produced by the redundant activity of several chromatin-modifying enzymes^10^. Therefore, attaining full depletion of particular histone methylations *in vivo* usually requires complex mouse genetic models that consist of multiple mutated genes in specific cell types^11^. Furthermore, chromatin-modifying enzymes have non-histone methylation substrates or methylation-independent activities^12–14^, thereby complicating the interpretation of the results using loss-of-function alleles for histone methyltransferases.

Somatic mutations that substitute a lysine (K) with methionine (M) at specific positions in the N-terminal tails of histone H3 were initially discovered in some types of cancer in humans^15^. Histone H3 K-to-M mutations function as dominant negative molecules by repressing the activity of the histone methyltransferases targeting a specific lysine residue on the endogenous histone H3, thereby blocking the methylation of that residue (Fig. 1)^16^. For example, experimental overexpression of a mutant histone H3 carrying a lysine 27-to-methionine substitution (H3K27M) acts primarily as an inhibitor of the H3K27 methyltransferase EZH2, resulting in the depletion of H3K27me3 in the endogenous histone H3^16^. Likewise, overexpression of histone H3K9M blocks the activity of H3K9 methyltransferases, such as SUV39H1 and G9a, leading to H3K9me3 depletion in the endogenous histone H3^16^. Moreover, integration of the histone K-to-M mutant into nucleosomes is expected to locally reduce H3 methylation, as methionine is not a substrate for methylation (Fig. 1)^17^.

**Fig. 1.**
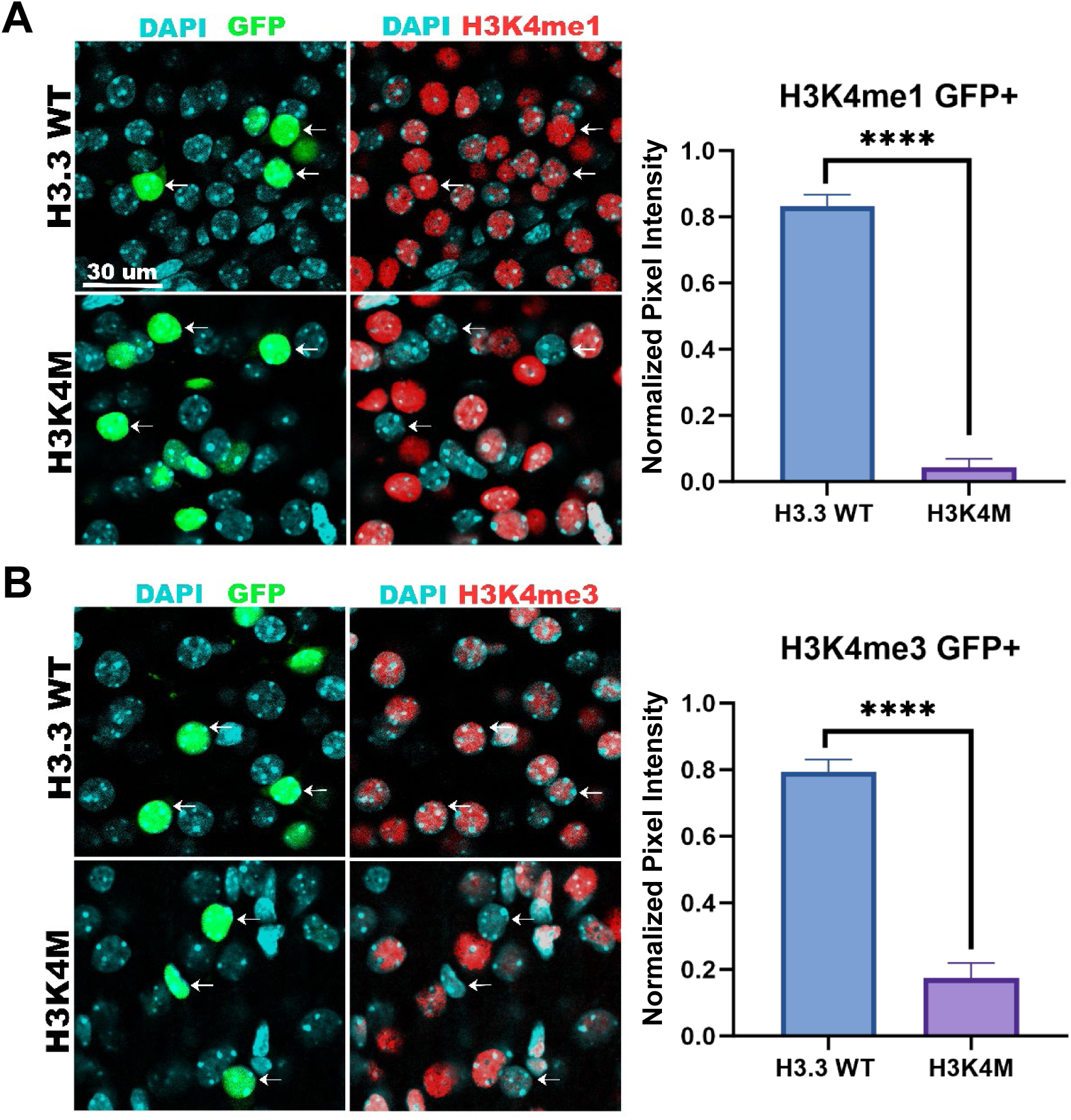
Inhibition of histone H3 methylation. Overexpression of K-to-M mutant histone H3 (yellow circle) results in the inhibition of specific groups of lysine methyltransferases, leading to the indirect depletion of lysine methylation (shown as “me” in red) on the endogenous histone H3 (indirect mechanism). The mutant transgene is also incorporated into the nucleosomes, which results in an unmethylated methionine (direct mechanism).

Several groups have implemented the experimental overexpression of histone H3 K-to-M mutants in multiple systems to deplete the endogenous histone H3 methylation without introducing mutations in the genes encoding for histone H3 methyltransferases^15^. This approach has been used in invertebrates to test the role of histone H3K27 methylation in Caenorhabditis elegans^18^ and H3K9, H3K27, and H3K36 methylation in Drosophila melanogaster^19–21^. In mammals, histone K-to-M mutants have been used to study the role of H3K9 and H3K36 methylation in mouse embryonic stem cells (mESCs) and hematopoietic cells *in vivo*^22^, as well as the function of H3K27 methylation in pediatric gliomas using human neural progenitors *in vitro*^23^. These studies indicate that histone H3 K-to-M mutants are effective tools to study chromatin biology in model organisms.

Here, we leveraged the use of histone K-to-M mutants to deplete various histone H3 methylations in developing and mature neurons of the mouse cerebral cortex. We present a set of molecular vectors to achieve three goals *in vivo*: 1) Robust and simultaneous depletion of two levels of histone H3 methylation at a single lysine residue; 2) Concurrent removal of histone H3 methylation at two independent lysine residues; 3) Temporal control of methylation depletion at specific stages of neuronal differentiation. Our tools can be implemented in other systems to study the regulation of gene expression by multiple levels of histone methylations and their crosstalk in specific cell types of developing and adult mice or *in vitro* human models of cell differentiation.

## RESULTS

We aimed at depleting specific histone H3 methylations in early postnatal neurons of the mouse cerebral cortex by performing *in utero* electroporation (IUE) of plasmids encoding for histone H3 K-to-M mutants. One caveat of this approach is that plasmids are initially electroporated into proliferating neural stem cells (NSCs) and progenitors at the ventricular zone of the embryonic cortex^24^, which is expected to result in reduced concentration of the molecular vectors over multiple rounds of cell division and lower gene expression levels upon neuronal differentiation. To decrease plasmid dilution in proliferating cells, we generated a series of Enhanced Episomal Vectors (EEVs) based on Epstein-Barr virus sequences^25^. The vectors consisted of a constitutively active CAG promoter driving the expression of the WT histone H3.3A isoform (*H3f3a*)^26^ or K-to-M mutants on lysine 4 (H3K4M), 9 (H3K9M), and 27 (H3K27M) upstream an internal ribosome entry site (IRES) and nuclear GFP (Supplementary Fig.1). The EEVs also contained the EBNA1 (Epstein–Barr virus nuclear antigen 1) and oriP (origin of replication) sequences (Supplementary Fig.1)^25^, which are known to maintain high transgene expression in proliferating mouse cells, possibly by enhancing nuclear retention of the molecular vector^27–30^.

We performed IUE of the plasmid encoding for WT histone H3.3 or H3K4M into embryonic day 14.5 (E14.5) mouse cerebral cortex, a developmental stage at which upper-layer (i.e. layers II-IV) excitatory neurons are born^31^. We collected the electroporated brains at postnatal day 8 (P8), a time point when upper-layer cortical neurons have completed their migration^31^. Our analysis focused on the depletion of H3K4 monomethylation (H3K4me1) and H3K4 trimethylation (H3K4me3). Using immunostaining, we quantified the levels of H3K4me1 and H3K4me3 in electroporated neurons, which also expressed nuclear GFP. Our analysis shows that H3K4M overexpression resulted in a robust and simultaneous depletion of H3K4me1 and H3K4me3 in comparison with the overexpression of the WT histone H3.3 (Fig. 2A, B). Most H3K4M-electroporated neurons expressed RBFOX3 (also known as NeuN) —a marker of mature neurons at P8 (Supplementary Fig. 2A). The electroporated neurons also expressed SATB2, a marker of callosal projections neurons in the cortex (Supplementary Fig. 2A)^32^. These results suggest that depletion of H3K4me1 and H3K4me3 does not lead to a general silencing of actively transcribed genes, as H3K4M-expressing cortical neurons survived and retained some molecular identity markers. Moreover, we combined the IUE of histone H3K4M with the supernova system^33^, which enabled the sparse labeling of electroporated neurons and the simultaneous depletion of H3K4me1 in comparison with the WT histone H3.3-electroporated controls (Supplementary Fig. 2B). This implementation will facilitate future studies directed at testing the role of histone methylations in the development of dendrites, axons, and synaptic contacts in cortical neurons.

**Fig. 2.**
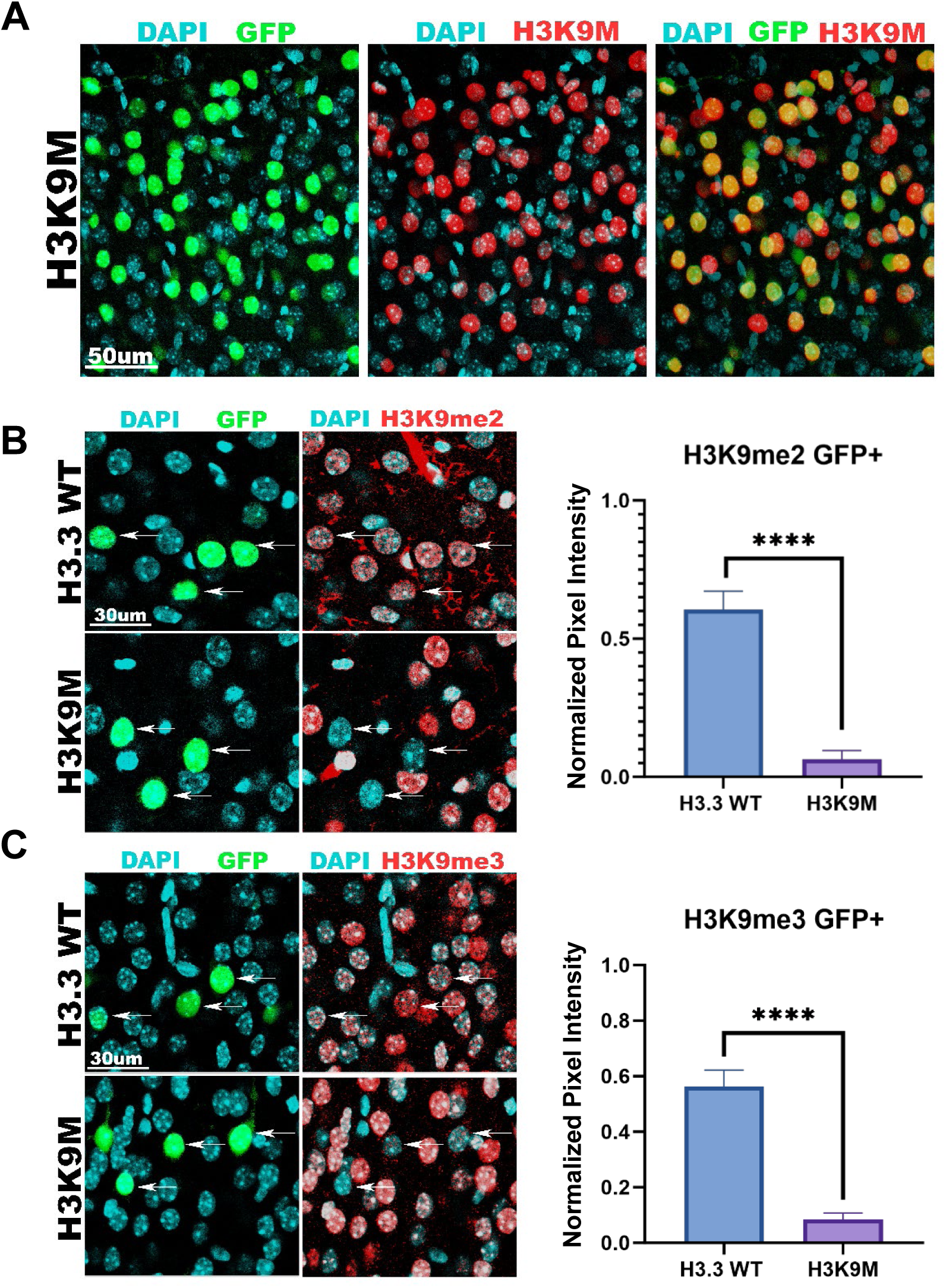
Depletion of Histone H3K4 methylation in cortical neurons. E14.5 cortices were electroporated *in utero* with EEVs carrying GFP and histone H3.3 WT or H3K4M. GFP^+^ cortical neurons were analyzed at P8. H3K4M produced a strong depletion of H3K4me1 (A) and H3K4me3 (B) in comparison to H3.3 WT electroporated controls. Arrows indicate some representative examples in each condition. Data represent mean ± SD; statistical analysis is unpaired Student’s t-test (****p < 0.001).

We sought to determine whether the EEV encoding histone H3K9M enabled the simultaneous depletion of the repressive marks H3K9me2 and H3K9me3 in cortical neurons. We performed IUE of EEV-H3K9M at E14.5, which resulted in strong expression of the mutant histone in all electroporated neurons at P8 (Fig. 3A). We observed a strong depletion of H3K9me2 and H3K9me3 in most GFP^+^ neurons electroporated with H3K9M in comparison to neurons electroporated with WT histone H3.3 at P7-P8 (Fig. 3B, C). In contrast, H3K9M expression did not have an overt impact on H3K9me1 —a histone mark associated with transcriptionally active genes (Supplementary Fig. 3A). Our findings indicate that expression of H3K9M leads to simultaneous depletion of the repressive marks H3K9me2 and H3K9me3 *in vivo*. This effect seems specific, as we did not find an apparent decrease of H3K27me3 in H3K9M-electroporated neurons at P8 (Supplementary Fig. 3B).

**Fig. 3.**
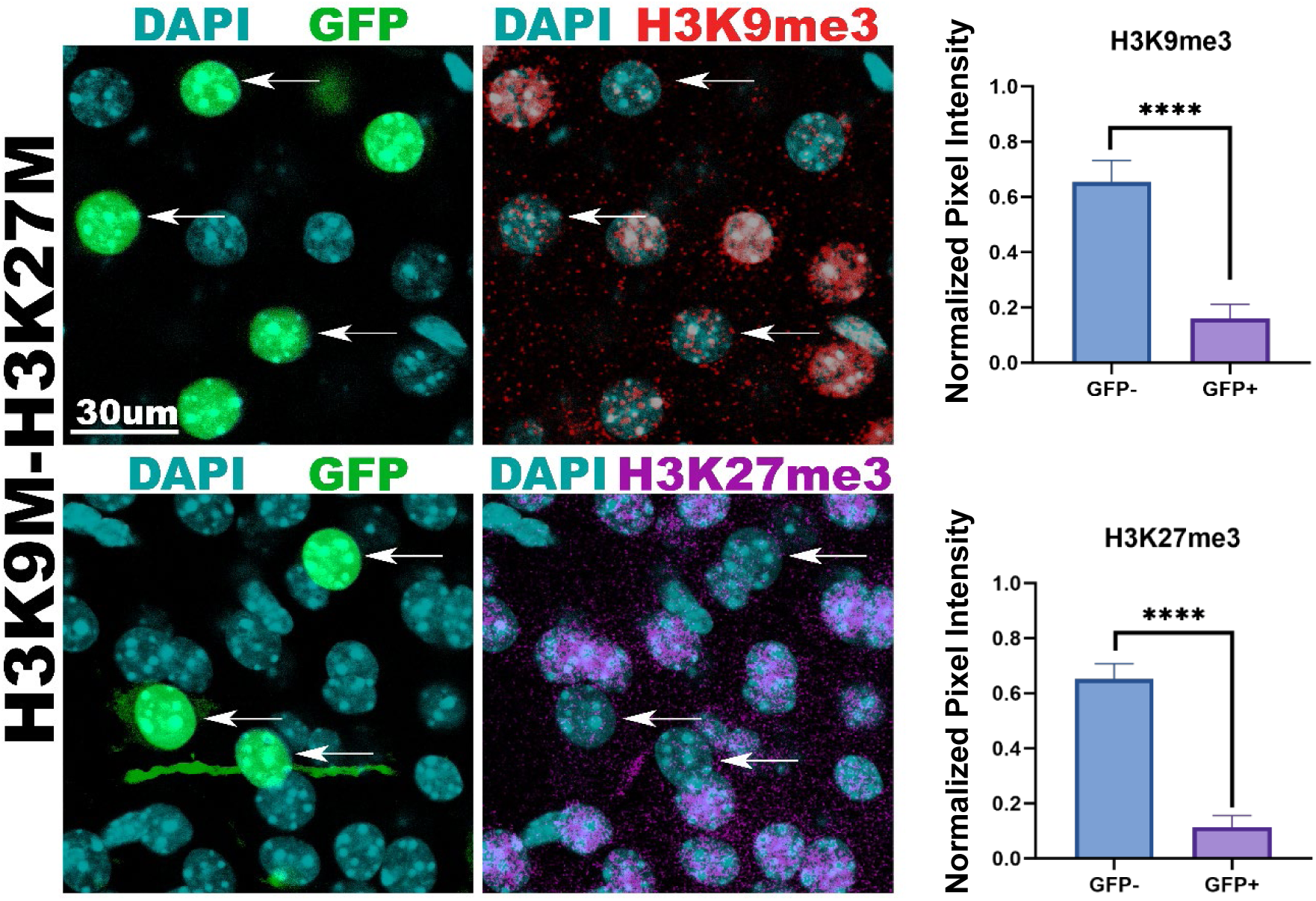
Depletion of Histone H3K9 methylation in cortical neurons. E14.5 cortices were electroporated *in utero* with EEVs carrying GFP and histone H3.3 WT or H3K9M. GFP^+^ cortical neurons were analyzed at P7-P8. High levels of H3K9M expression were confirmed in all electroporated neurons (A). H3K9M produced a strong depletion of H3K9me2 (B) and H3K9me3 (C) in comparison to the H3.3 WT electroporated controls. Arrows indicate some examples. Data represent mean ± SD; statistical analysis is unpaired Student’s t-test (****p < 0.001).

To test whether the expression of mutant histones facilitates the simultaneous depletion of H3K9me3 and H3K27me3 in cortical neurons, we performed IUE of EEVs encoding H3K9M and H3K27M at E14.5 followed by the analysis of histone marks at P15. We observed a strong and simultaneous depletion of H3K9me3 and H3K27me3 in electroporated neurons in comparison to the non-electroporated cells (Fig. 4). In this experimental setting, H3K9me2 was depleted in some electroporated neurons whereas others retained the histone mark (Supplementary Fig. 4), likely as a consequence that the concentration of the two injected plasmids (H3K9M and H3K27M) was around half the concentration of the individually injected H3K9M (see the Methods section). Therefore, the robust depletion of H3K9me2 and H3K9me3 described above (Fig. 3B, C) likely depends on the amount of H3K9M that is expressed in developing neurons. Likewise, the expression of H3K27M produced a decrease of H3K27me2 in a subset of electroporated neurons (Supplementary Fig. 4). Together, our results suggest that this experimental approach is suitable for elucidating the functional interplay of H3K9 and H3K27 methylation in transcriptional repression and cell differentiation.

**Fig. 4.**
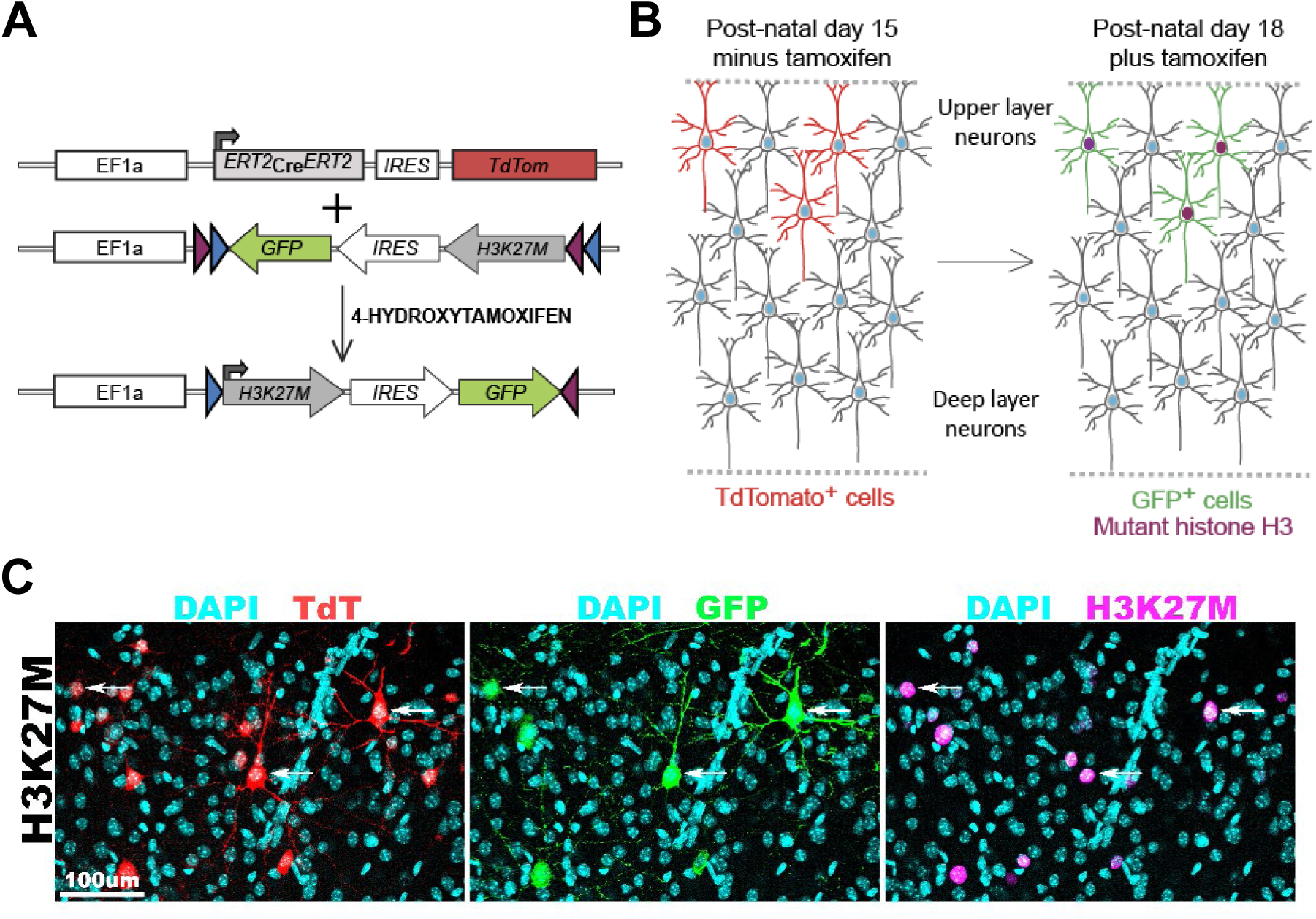
Simultaneous depletion of Histone H3K9 and H3K27 methylation in cortical neurons. E14.5 cortices were electroporated *in utero* with EEVs carrying GFP, H3K9M, and H3K27M. GFP^+^ cortical neurons were analyzed at P15. A significant decrease of H3K9me3 and H3K27me3 was observed in electroporated cortical neurons (arrows) in comparison to nonelectroporated GFP^-^ cells. Data represent mean ± SD; statistical analysis is unpaired Student’s t-test (****p < 0.001).

To control the timing of mutant histone expression in cortical neurons, we generated a conditional system consisting of two vectors based on Cre-LoxP recombination. The first vector encodes a tamoxifen-inducible Cre that is flanked by the estrogen receptor ligand-binding domain (ER^T2^) on the N- and C-terminal ends (Fig. 5A). The inducible Cre is produced in a bicistronic transcript with tdTomato to label electroporated neurons independently of tamoxifen administration (Fig. 5A, B). The second vector contains a FLEX (Flip-Excision) switch that is irreversibly flipped upon Cre-mediated recombination of the LoxP/Lox2272 flanking sites to initiate expression of the WT or mutant histone H3 and GFP (Fig. 5A, B). To test for potential leakage of Cre activity, we initially injected both vectors at E14.5 and collected samples at P15 in the absence of tamoxifen administration. We found that the electroporated cortical neurons were labeled by tdTomato but no detectable levels of GFP were observed at the time of analysis (Supplementary Fig. 5), confirming that the inducible Cre that is flanked by ER^T2^ on the N- and C-terminal ends has no leakage^34^. Next, we performed IUE at E14.5 of the Cre-inducible system using a FLEX vector coding for H3K27M, followed by tamoxifen administration in electroporated pups from P15 to P18. We observed the induction of GFP and H3K27M in all electroporated neurons at P18 (Fig. 5C). This result indicates that the inducible system enables the temporal control of gene induction in the cerebral cortex.

**Fig. 5.**
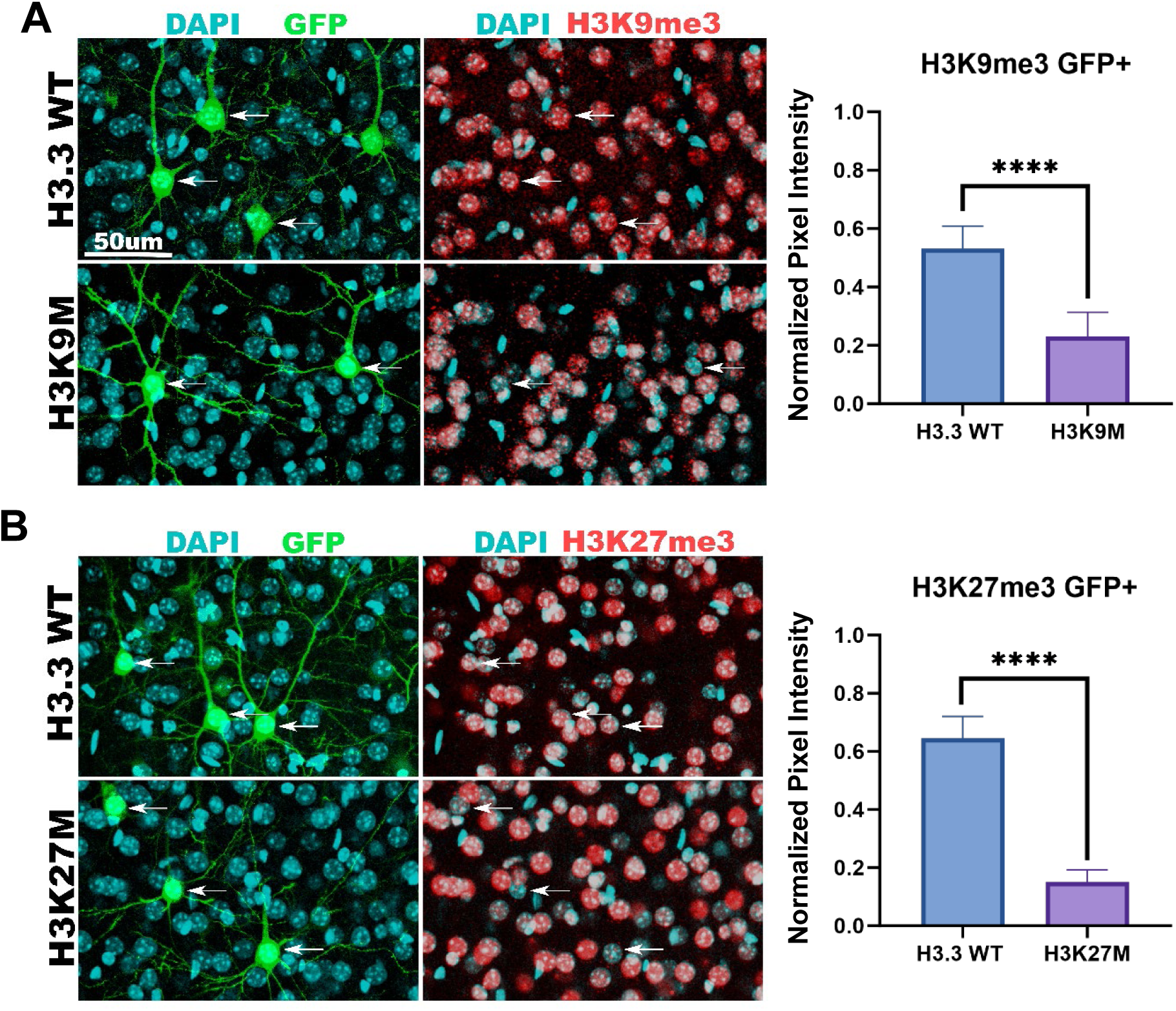
A conditional vector system for the temporal control of histone H3 expression. (A) The schematic shows a vector consisting of a tamoxifen-inducible Cre and tdTomato, and a second vector with a FLEX cassette carrying H3K27M and GFP. (B) Experimental strategy to activate H3K27M in electroporated neurons of the postnatal cortex. (C) E14.5 cortices were electroporated with the two vectors shown in (A) and 4-hydroxytamoxifen was administered from P15 to P18. The electroporated cortical neurons displayed expression of TdTomato, GFP, and H3K27M at P18 (arrows).

To test whether mutant histones promote the depletion of endogenous histone marks in the postnatal cortex, we performed IUE of WT histone H3.3, H3K27M, or H3K9M at E14.5, followed by tamoxifen administration in electroporated pups at P15-P18, and the analysis of histone marks at P70. We observed that the WT histone H3.3 did not impact H3K9me3 or H3K27me3 (Fig. 6A, B). In contrast, H3K9M and H3K27M induced the depletion of H3K9me3 and H3K27me3, respectively (Fig. 6A, B). Using this experimental design, we also observed depletion of H3K9me3 and H3K27me3 in electroporated neurons as late as P100 (Supplementary Fig. 6A, B), indicating that inhibition of histone methylation is sustained in postmitotic cells for at least 2.5 months after tamoxifen administration. Our results show that the tamoxifen-inducible vector system enables the depletion of specific histone H3 methylations in fully differentiated cortical neurons in the early postnatal and adult brains.

**Fig. 6.**
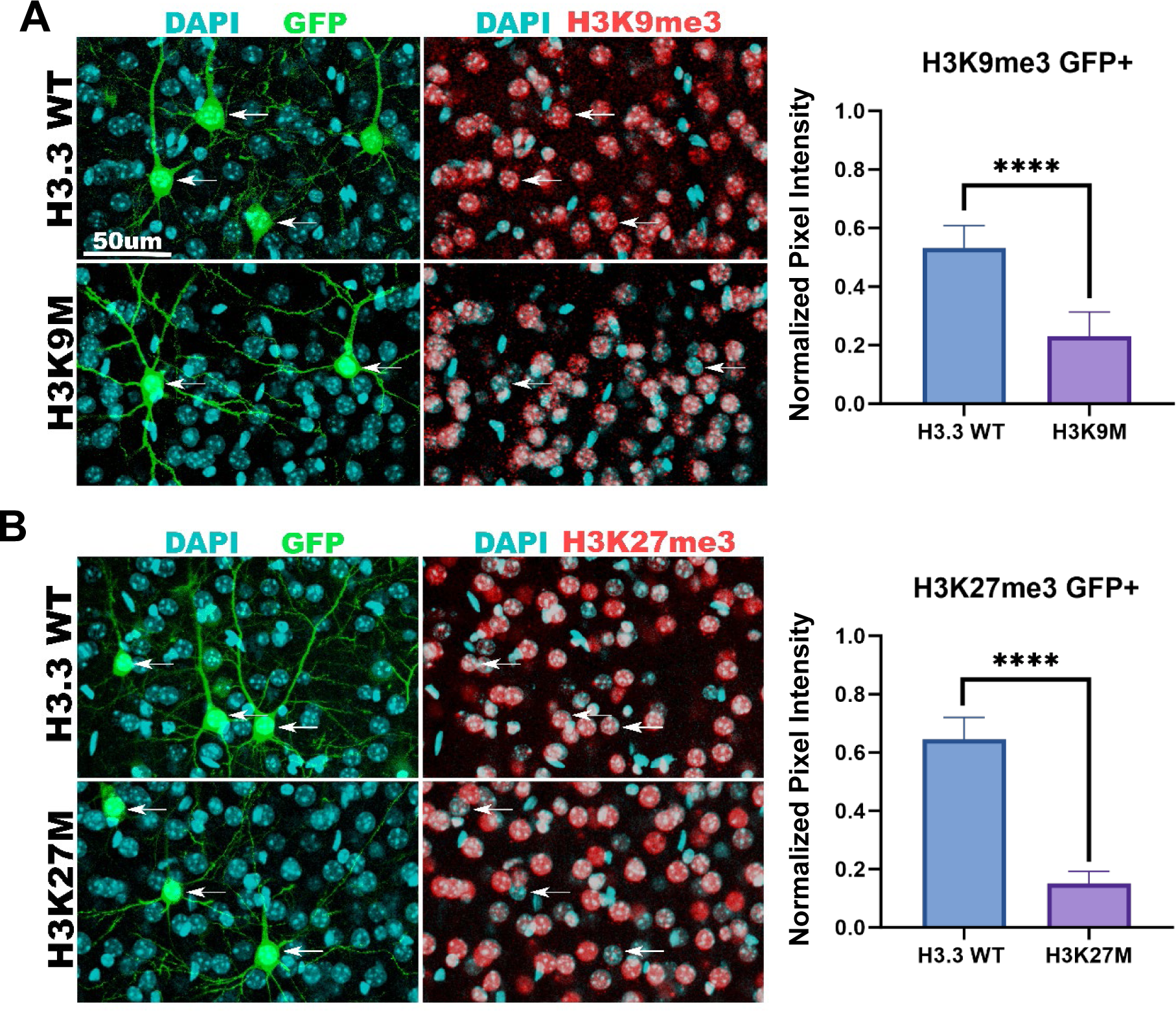
Inducible depletion of histone H3K9 or H3K27 methylation in cortical neurons. E14.5 cortices were electroporated *in utero* with vectors carrying the inducible Cre and FLEX cassette, followed by tamoxifen administration from P15 to P18 to activate histone H3 genes. Histone H3 methylation was quantified in GFP^+^ cortical neurons at P70. H3K9M produced a significant decrease of H3K9me3 in comparison to histone H3.3 WT (A, arrows), whereas H3K27M expression resulted in depletion of H3K27me3 (B, arrows). Data represent mean ± SD; statistical analysis is unpaired Student’s t-test (****p < 0.001).

## DISCUSSION

Elucidating the role of genome-wide histone H3 methylations in transcriptional regulation and cell differentiation remains a major challenge in mammals^4,8^. One significant hurdle is the presence of multiple copies of histone H3-coding genes in mammalian genomes. In mice for example, histone H3 has three isoforms (H3.1, H3.2, H3.3) encoded by 28 alleles^35^. Notably, a recent study using CRISPR-mediated gene editing in cultured mESCs simultaneously mutated the sequence encoding lysine 27 in all histone H3 alleles^36^. However, implementing this type of approach in a cell-type-specific manner *in vivo* remains a daunting technical challenge in mice. Furthermore, multiple histone H3 methyltransferases normally play redundant roles in the methylation of specific lysine residues^10^. Therefore, conditional mutations of individual methyltransferases do not usually result in a complete loss of histone H3 methylation^12,37–39^. To overcome these limitations, we have optimized the overexpression and temporal control of histone H3 K-to-M mutants to inhibit the endogenous methylation of histone H3 in early postnatal and adult cortical neurons. We demonstrate that this relatively simple approach enables the robust depletion of at least two levels of methylation at individual or multiple lysine residues of histone H3 *in vivo*.

Previous studies have shown that histone H3 K-to-M mutants can inhibit methylation on a particular lysine residue with different strengths. H3K27M and H3K36M are strong inhibitors of the di- and trimethylated states of lysine 27 and 36, respectively^16,22^, whereas H3K4M and H3K9M are weaker at inhibiting multiple methylation states^22,40^. In this regard, previous work showed that H3K4M overexpression in mouse adipocytes strongly silences H3K4me1 but has a mild effect on H3K4me3^40^. Although H3K4M expression was driven under a strong CAG promoter^40^, the transgene integration into the host chromatin environment might reduce H3K4M levels and H3K4me3 depletion in adipocytes. In another study, H3K9M overexpression in mouse hematopoietic cells inhibited H3K9me3 but had little impact on H3K9me2, possibly as a consequence of H3K9M integration into the host genome as a single copy^22^. Here, we show that sustained high expression of H3K4M or H3K9M leads to a strong depletion of H3K4me1/3 or H3K9me2/3, respectively, in cortical neurons.

In our experimental setting, the initially electroporated cells in the E14.5 cerebral cortex primarily consist of proliferating NSCs and progenitors residing at the ventricular zone^24^. Standard molecular vectors are diluted more rapidly than EEVs during mitosis, resulting in decreased gene expression in proliferating cells and their differentiated progeny^27^. EEVs display nuclear retention and self-replicate as extrachromosomal elements in human cells, thereby maintaining high gene expression levels upon multiple rounds of cell division^25,41^. EEVs do not self-replicate in mouse cells but display high nuclear retention during mitosis^27–30^. Nuclear retention is mediated by the binding of EBNA1 protein to the direct repeat element of oriP^42^ —a mechanism that promotes the attachment between the plasmid and the nuclear matrix chromosomal scaffold in mitotic cells^43^. Consistent with this notion, our results indicate that EEVs maintain high expression of mutant histones during proliferation and neuronal differentiation, resulting in robust depletion of two levels of histone H3 methylation.

We observed that a high concentration of EEV-H3K9M robustly inhibited H3K9me2 and H3K9me3, whereas a medium concentration of EEV-H3K9M strongly depleted H3K9me3 but had a partial impact on depleting H3K9me2. Distinct canonical histone methyltransferases are responsible for generating different H3K9 methylation states —EHMT1 (GLP) and EHMT2 (G9A) produce H3K9me2, while SETDB1, SETDB2, SUV39H1, and SUV39H2 generate H3K9me3^8,44^. Previous evidence indicates that H3K9M can inhibit EHMT2 and SUV39H1 *in vitro*^16^. However, our evidence suggests that H3K9M is less effective *in vivo* at inhibiting EHMT1/2 than at repressing SETDB1/2 and SUV39H1/2, as a higher concentration of H3K9M seems necessary to suppress EHMT1/2 activity. Unexpectedly, we did not observe depletion of H3K9me1 —a mark also generated by EHMT1/2^8,44^. One possibility is that noncanonical H3K9 methyltransferases that are not sensitive to H3K9M-mediated inhibition might be compensating for the partial loss of EHMT1/2 activity in cortical cells. In this regard, it is interesting that the genes encoding for the noncanonical H3K9 methyltransferases PRDM16 and PRDM8 are highly expressed in NSCs and neurons of the embryonic cortex, respectively^45–48^. Further studies are necessary to determine the full range of canonical and noncanonical histone methyltransferases that are repressed by mutant histones and their differential sensitivities to inhibition.

An outstanding open question in the field of epigenetics is how the coordinated activities of H3K9me3 and H3K27me3 regulate developmental gene expression and cell fate^9^. H3K9me3 and H3K27me3 are enriched within domains of constitutive and facultative heterochromatin^49^. Although H3K9me3- and H3K27me3-decorated heterochromatin display only a partial overlap, the loss of H3K9me3 leads to the expansion of H3K27me3 and vice versa^49–53^. These observations suggest that H3K9me3 and H3K27me3 may cooperate in silencing transcription. However, the simultaneous depletion of H3K9me3 and H3K27me3 using conventional mouse genetics entails the generation of a line carrying at least six conditional loss-of-function mutations in the genes coding for the canonical H3K9me3 methyltransferases (*Setdb1*, *Setdb2*, *Suv39h1*, and *Suv39h2*) and H3K27me3 methyltransferases (*Ezh1* and *Ezh2*) —an unpractical approach. To overcome this limitation, we have leveraged the use of histone H3K9M and H3K27M mutants to concomitantly block the activity of H3K9 and H3K27 methyltransferases. Our results show a substantial and simultaneous depletion of H3K9me3 and H3K27me3 in cortical neurons. Future experiments with the vectors described in this study may be directed at testing the functional interplay of H3K9me3 and H3K27me3 in heterochromatin formation and the silencing of lineage-inappropriate genes in the developing cerebral cortex.

We report the generation of a conditional Cre-FLEX system to control the timing of depletion of specific histone H3 methylations. In comparison to the EEVs described here, the FLEX vectors offer the advantage of activating the mutant histones specifically in post-natal cortical neurons. Our results indicate that the decrease of H3K27me3 and H3K9me3 using the FLEX system is significant, albeit not as robust as the depletion observed with the EEVs vectors. This difference suggests that histone H3 mutants are more effective in decreasing histone methylation during the early stages of neuronal differentiation than in mature neurons. This observation might reflect the fact that neuronal differentiation is accompanied by global changes in chromatin modifications^54^. Supporting this idea, recent findings indicate that accumulation of the histone H3.3 isoform in differentiating cortical neurons is important to establish their identity and axonal connectivity^55^. Therefore, the induction and incorporation into nucleosomes of histone H3.3 during cortical neurogenesis could favor the generation of methylation-depleted histones in the presence of K-to-M mutants. In contrast, a consolidated chromatin landscape in mature neurons, together with a relatively slow histone H3 turnover^56^, could reduce the generation rate of methylation-depleted histones. We propose that the combination of the EEVs and FLEX vectors described here will facilitate the distinction between the role of histone methylation in developmental lineage specification and mature cell functions.

Loss-of-function mutations in histone H3 methyltransferases and genome-wide changes in histone H3 methylation are associated with multiple neurodevelopmental disorders, brain tumors, aging, and neurodegeneration^57–59^. The experimental use of histone H3 K-to-M mutants in invertebrate and vertebrate model organisms will contribute to our understanding of the role of histone H3 methylation in various diseases. One important limitation of this approach is that the mechanism by which histone K-to-M mutants inhibit the endogenous methylation levels is not completely understood^15^. Future experiments must be directed at comparing mutant histone overexpression with other approaches, such as genetic or pharmacological models targeting specific histone methyltransferases. Nevertheless, we provide evidence that mutant histone overexpression is a relatively simple starting point to test the combined role of at least two levels of methylation at individual or distinct lysine residues of histone H3. Moreover, the protein sequences of mouse and human histone H3.3A are identical, suggesting that the vectors presented here should also inhibit histone methylation in human cells. Another advantage is that EEVs self-replicate in human cells, offering the possibility of long-term expression without genomic integration^41^. This approach was used to reprogram adult fibroblasts into human-induced pluripotent stem cells by transfection of EEVs expressing pluripotency factors^60^. Implementation of the tools described herein in mouse and human cell lineages that are easily accessible by electroporation or transfection will expand the use of mutant histones in exploring the role of the epigenome in development and disease.

## METHODS

### Experimental model

All the animal procedures conducted in this study complied with the protocol approved by the Institutional Animal Care and Use Committee (IACUC) of Indiana University. All mice used in this study were ICR (CD-1) outbred from Envigo. Embryonic (E) day 14.5 and postnatal (P) day 7-70 mice were used in this study. The sex of the mouse embryos and pups was not determined. Mouse housing, husbandry, medical treatment, and euthanasia procedures followed the standards set by the Laboratory Animal Resources at Indiana University Bloomington. The authors complied with the ARRIVE guidelines.

### *In utero* electroporation and tamoxifen administration

Timed pregnant mice were anesthetized by isoflurane inhalation (3-5% for induction, 1-3% for maintenance). The anesthetized dams received a subcutaneous dose of buprenorphine (0.05-0.1 mg/kg) and were placed on top of a warming pad. The abdominal hair was removed using an electric clipper and the shaved abdomen was cleaned three times by alternating Iodine and alcohol wipes. An abdominal midline incision was made and the uterine horns were exposed on top of a sterile gauze pad. During the entire procedure, the open peritoneal cavity and the exposed uterine horns were kept moist with PBS 1X at 37° C. All electroporated plasmids were purified with an endotoxin-free maxi kit (Invitrogen) and dissolved in sterile PBS 1X containing 0.025% Fast Green (SIGMA). Approximately 1 μl of plasmid solution was manually injected into the forebrain ventricle of each embryo using heat-pulled/beveled glass micropipettes (Drummond). Once the whole litter was injected, 5 pulses of 30-40 V (50 ms duration and 950 ms intervals) were applied with 7 mm platinum electrodes connected to an ECM 830 square wave electroporator (BTX). The uterine horns were re-introduced into the peritoneal cavity and the incision was closed with an absorbable suture on a curved needle using a continuous locking stitch. The skin was then closed with sterile stainless steel wound clips. Ketoprofen (5 mg/kg) was subcutaneously administered to the anesthetized mouse to reduce pain. Mice were allowed to recover in a chamber set at 37° C until they ambulated stably about the chamber. A second dose of ketoprofen was administered 24 hours post-surgery and the animals were monitored every day during the next seven days. The EEV encoding WT and mutant histone H3.3 were individually injected at 2-3 μg/μl. The Supernova plasmids pK031 (pTRE-Cre) and pK029 (pCAG-loxP-stop-loxP-RFP-IRES-tTA-WPRE)^33^ were injected at 1.0 and 0.5 μg/μl, respectively, in combination with 2.0 μg/μl of the EEV encoding H3K4M or WT H3.3. The EEVs carrying H3K9M and H3K27M were co-injected at 1.5 μg/μl each. For the inducible system, the FLEX vectors carrying WT H3.3, H3K9M, or H3K27M were injected at 2 μg/μl in combination with 1 μg/μl of the plasmid encoding the tamoxifen-inducible Cre. Tamoxifen (SIGMA, T5648) was dissolved in sterile corn oil (SIGMA, C8267) at 20 mg/ml and aliquots were frozen at −20° C to prevent variability between batches. For induction of Cre-mediated recombination of the FLEX cassettes, tamoxifen was administered subcutaneously at 8 mg/40 gm of body weight in mouse pups every day from P15 to P18, as previously described^61^.

### Cloning and site-directed mutagenesis

The mouse *H3f3a* sequence (encoding the histone H3.3A isoform) upstream an IRES and nuclear GFP sequences were cloned into the EcoRI and NotI sites of EEV600A-1 (System Biosciences) to generate the EEV expressing the WT histone H3.3A (supplementary Fig. 1). A Kozak consensus sequence (GCCACCATGG) in *H3f3a* was included in this vector to enhance the translation efficiency of histone H3.3A. Custom primers were designed to mutate lysine 4, 9, or 27 into methionine on histone H3.3A using the QuikChange II XL Site-Directed Mutagenesis Kit (Agilent Technologies #200521) following the manufacturer’s instructions. All mutations were sequence-confirmed using custom primers.

To generate the vectors carrying the inducible histone H3.3, we cloned an IRES-GFP sequence into the SpeI site of pJ241-FLEX (Addgene #18925). To drive expression of the FLEX cassette under the EF1a promoter, the FLEX [IRES-GFP] sequence was introduced into the EcoRV-KpnI sites of pAAV-EF1a-FLEX-GTB (Addgene #26197) using infusion cloning (Takara Bio). This step swapped the FLEX cassette of pAAV-EF1a-FLEX-GTB for the FLEX [IRES-GFP] cassette. Finally, the WT and mutants histone H3.3A were cloned into the EcoRI site of the FLEX [IRES-GFP] sequence to produce a plasmid in which the EF1a promoter controls expression of a FLEX [histone H3.3A-IRES-GFP] cassette that produces functional gene products upon Cre-mediated recombination (Fig. 5A). To generate a vector encoding a tamoxifen-inducible Cre, we cloned CreER^T2^ (Cre recombinase fused to a mutant ligand-binding domain of the estrogen receptor) and IRES into the BamHI site of pAAV-EF1a-tdTomato-WPRE-pGHpA (Addgene #67527). To prevent leakage of Cre activity^34^, we ligated a Kozak-ER^T2^ sequence upstream of the Cre to produce a vector encoding ER^T2^CreER^T2^-IRES-tdTomato downstream of the EF1a promoter. All vectors were sequence-confirmed.

### Immunostaining

Early post-natal (P7-P15) and adult (P70) mice were anesthetized with an intraperitoneal injection of Xylazine (20 mg/Kg) - Ketamine (120-150 mg/Kg) and their brains were crosslinked by transcardially perfusing cold PBS pH 7.4 followed by 4% ethanol-free paraformaldehyde (PFA) diluted in PBS. The perfused brains were dissected and incubated overnight in 4% PFA with rocking at 4° C. The brains were washed three times in PBS and sectioned (75-100 μm thickness) in a VT1000 Vibratome (Leica Biosystems). Tissue sections were incubated in antigen retrieval solution (10 mM sodium citrate, 0.05% Tween 20, pH 6.0) for 1 hr at 70° C, colded down at room temperature (RT), and washed 3 times with PBS. Tissue sections were incubated for 1 hr at RT in blocking buffer (10% goat serum, 0.1% Triton X-100, 0.01% sodium azide diluted in PBS) before adding primary antibodies diluted in blocking buffer. Sections in primary antibody solution were incubated overnight at 4° C, washed three times with PBS, and incubated for 2 hr at RT with Alexa Fluor-conjugated secondary antibodies diluted in blocking buffer. Sections were washed again and the nuclei were stained with DAPI (4’,6-diamidino-2-phenylindole) before mounting with Fluoromount-G (Southern Biotech). Primary antibodies against histone H3 methylation on lysine 4, 9, and 27 were selected based on The Histone Antibody Specificity Database (http://www.histoneantibodies.com/)^62^. We used the following primary antibodies and dilutions: anti-H3K4me3 (Abcam, Ab8580, 1:6000), anti-H3K4me1 (Cell Signaling Technology, 5326T, 1:1000), anti-H3K9me3 (Active Motif, 39062, 1;1000), anti-H3K9me2 (Abcam, Ab1220, 1:1000), anti-H3K9me1 (Abcam, Ab8896, 1:1000), anti-H3K27me3 (Cell Signaling Technology, 9733S, 1:1000), anti-H3K27me2 (Active Motif, 7408001, 1:1000), anti-H3K9M (Invitrogen, MA5-33389 1:1000), anti-H3K27M (Cell Signaling Technology, 74829S, 1:2000), anti-GFP (Aves Labs Inc, GFP-1020, 1:4000), anti-RFP (MBL, PM005, 1:500), anti-NeuN (Millipore, MAB377, 1:300), anti-SATB2 (Santa Cruz, Sc-81376, 1:200). To detect primary antibodies, we used anti-rabbit and anti-mouse secondary antibodies conjugated to Alexa Fluor 488, 546, or 647 (Invitrogen, 1:500).

### Imaging and quantifications

For each experimental condition, brains were collected from two CD-1 mouse litters. A minimum of 3 immunostained sections from two brains were used for imaging with a Leica SP8 confocal microscope and processed with the Las X Life Science microscope software (Leica Microsystems). Imaging settings were identical between the experimental and control groups. The pixel intensity of histone methylation was manually measured in 30-70 electroporated (GFP^+^) or non-electroporated (GFP^-^) cells for each condition using Image J/Fiji (National Institute of Health, USA). Briefly, single-plane images were converted to 16-bit files, and the threshold values of the GFP channel were adjusted to manually create regions of interest (ROIs) surrounding multiple GFP^+^ cells on an image. ROIs were then overlaid with the DAPI channel to ensure the selected cells did not overlap with other cells. For GFP^-^ cells, ROIs were created in the DAPI channel. ROIs were then overlapped with the histone mark channel set at the default signal threshold and the pixel intensity for the histone methylation signal was measured within multiple ROIs. To reduce bias, ROIs were exclusively defined on the GFP or DAPI channels and without examining the histone methylation channel. To normalize for differences in signal intensity, we used the formula X New = X – Xmin / Xmax – Xmin, in which X represents the pixel intensity of the histone methylation staining in a cell. Accordingly, Xmin and Xmax were defined as the minimum and maximum pixel intensities of the histone methylation in electroporated cells for both control and experimental groups. Significance was determined using a two-tailed unpaired Student’s t-test with the Prism Software (GraphPad) and reported as *p<0.05, **p< 0.01, ***p<0.001, ****p< 0.0001.

## Data availability

The molecular vectors described in this paper are available from the corresponding author upon request.

## Author’s contributions

S.W., S.X., and D.R. performed all the experiments. S.W. quantified the results and prepared the figures. J-M.B. conceived the project, designed the experiments, and wrote the manuscript. All authors read and approved the final version of the manuscript.

