## Supplemental Figures for "A Vector System Encoding Histone H3 Mutants Facilitates Manipulations of the Neuronal Epigenome"

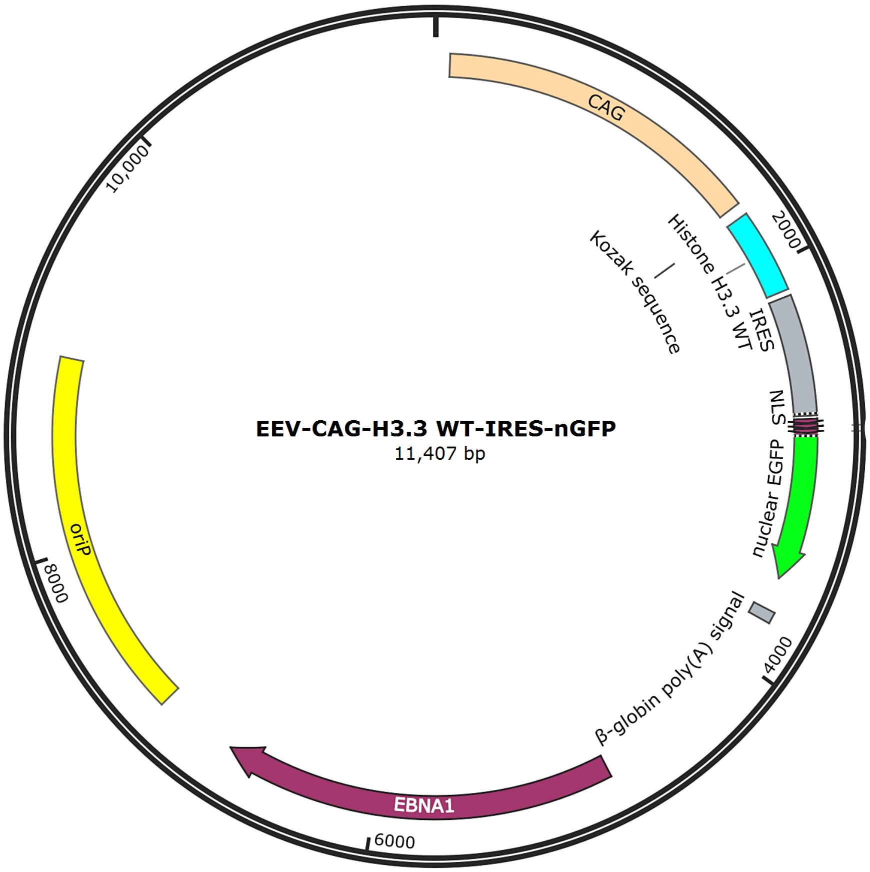


**Fig S1. Enhanced Episomal Vector (EEV) Map.** A series of EEVs were generated that encode the histone H3.3 WT (shown in the SnapGene vector map), H3K4M, H3K9M, or H3K27M upstream of an IRES (Internal Ribosome Entry Site) and nuclear GFP. The CMV early enhancer/chicken β actin (CAG) promoter drives the expression of histone H3 and GFP. NLS: Nuclear Localization Signal. EEVs contain the oriP and EBNA1 sequences that enhance nuclear retention of the plasmid in proliferating mouse and human cells and promote self-replication of the vector in dividing human cells.

.

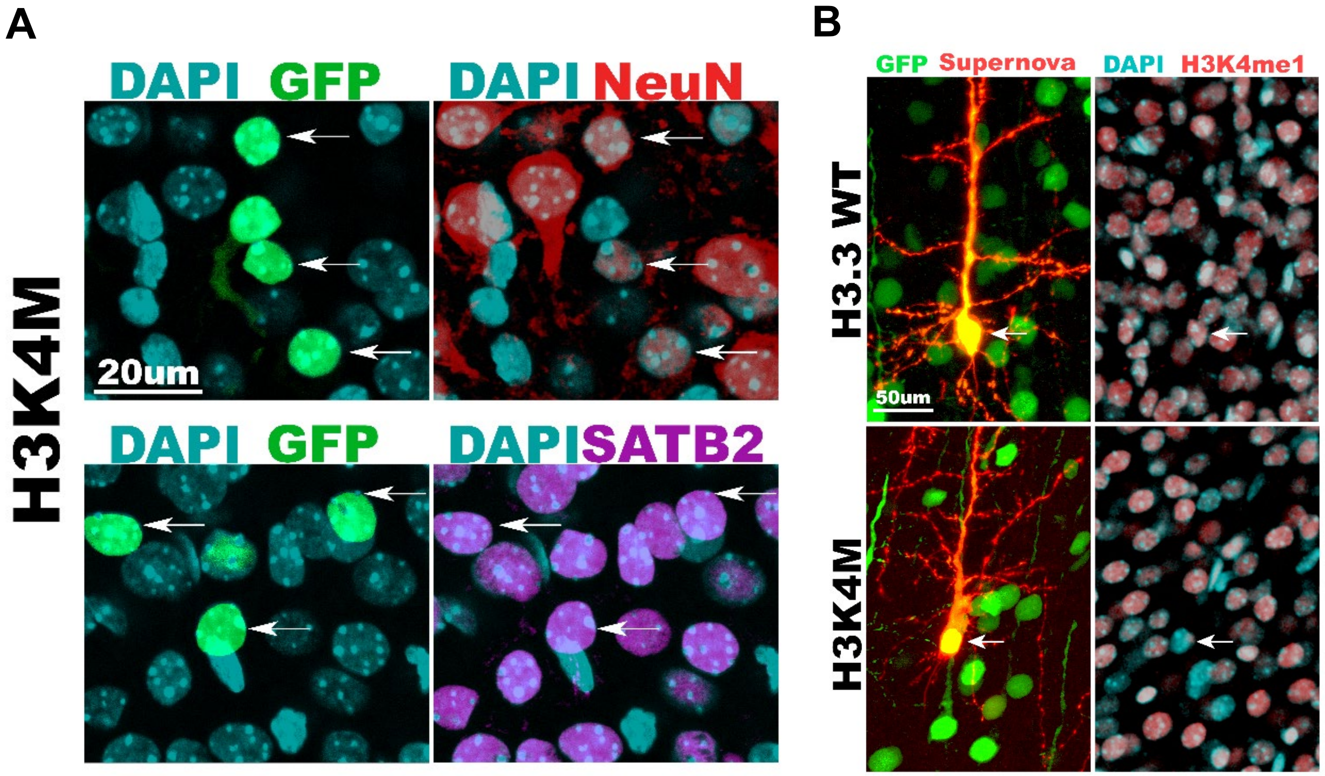


**Fig S2. Identity markers and morphology of H3K4M-electroporated cortical neurons.** (A) The majority of cortical neurons electroporated with H3K4M displayed expression of the mature neuron marker NeuN and the callosal projection neuron marker SATB2 at P8 (arrows). (B) Expression of H3.3 WT or H3K4M was combined with the Supernova system *in vivo* to sparsely label the morphology of some $GFP^{+}$ neurons with the red fluorescent protein (RFP, indicated as "Supernova"). The images on the left (GFP and RFP signals) are maximum projections of Z-stack images, whereas the images on the right (DAPI and H3K4me1) are single optical planes. Histone H3K4M depleted H3K4me1 in comparison to H3.3 WT at P8 (arrows).

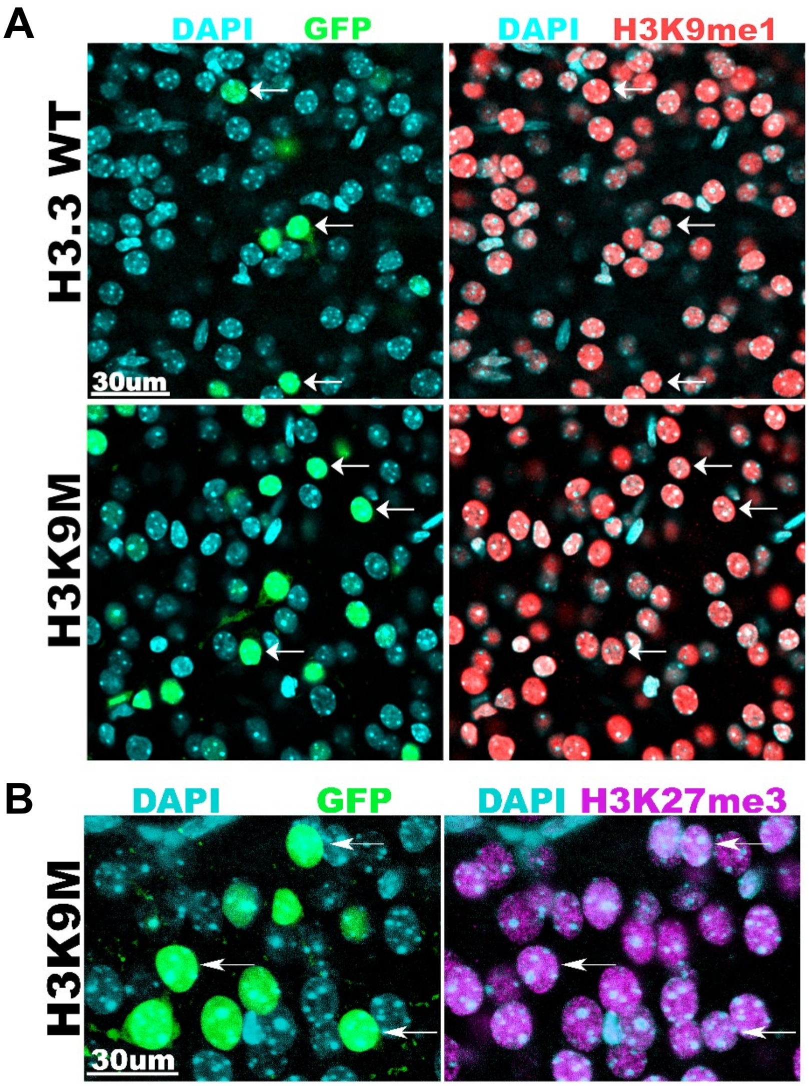


**Fig S3. Impact of H3K9M on histone methylation.** (A) Overexpression of H3K9M did not deplete H3K9me1 in GFP$^{+}$ cortical neurons at P8 (arrows). (B) H3K27me3 was not overtly reduced at P8 upon overexpression of H3K9M in cortical neurons (arrows).

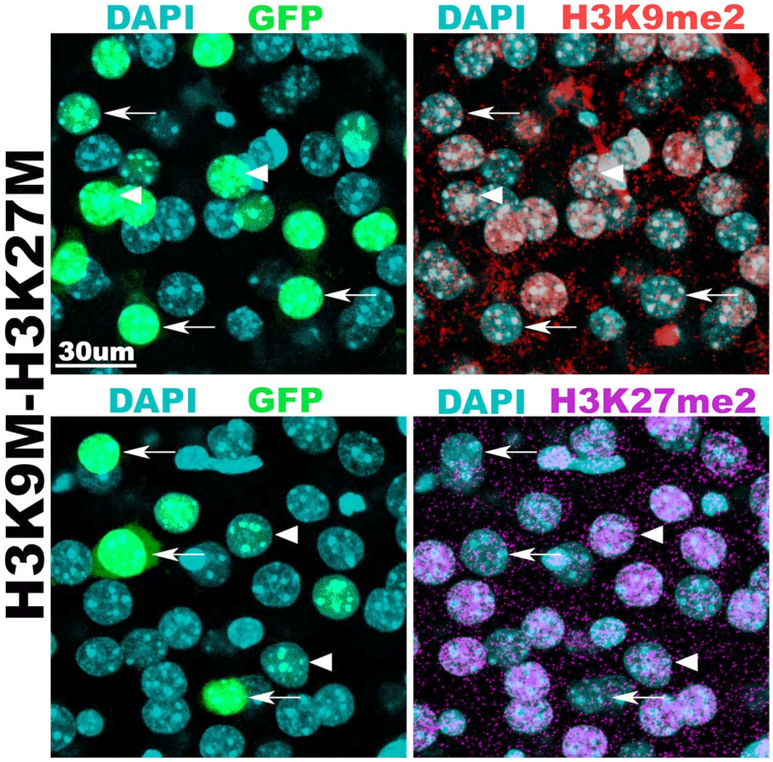


**Fig S4. Impact of H3K9M and H3K27M coexpression on H3K9 and H3K27 dimethylation.** Some cortical neurons simultaneously electroporated with H3K9M and H3K27M displayed depletion of H3K9me2 and H3K27me2 (arrows), whereas a different subset of electroporated neurons did not show an apparent decrease of the histone marks (arrowheads).

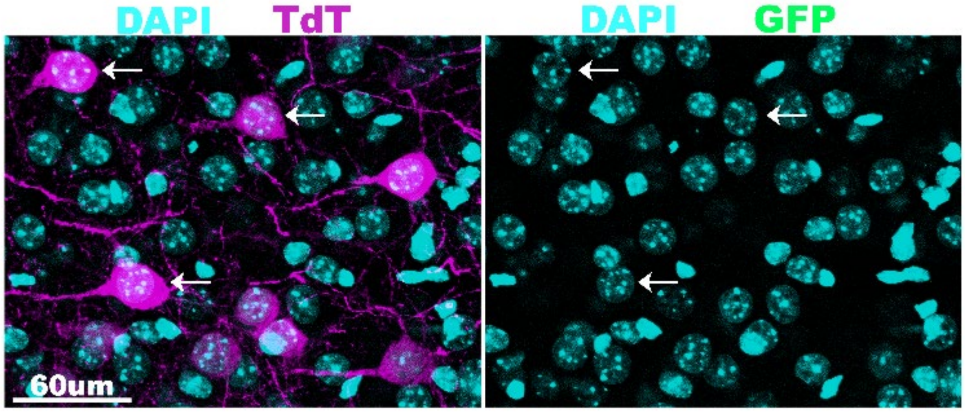


**Fig S5. No leakage of Cre activity.** The inducible Cre/FLEX vector system (see Fig. 5A) was electroporated at E14.5 and cortical neurons were analyzed at P15. TdTomato$^{+}$ cells (TdT) did not display detectable levels of GFP by immunostaining (arrows show some examples) indicating that the inducible $ER^{T2}CreER^{T2}$ does not promote recombination of the FLEX cassette in the absence of tamoxifen administration.

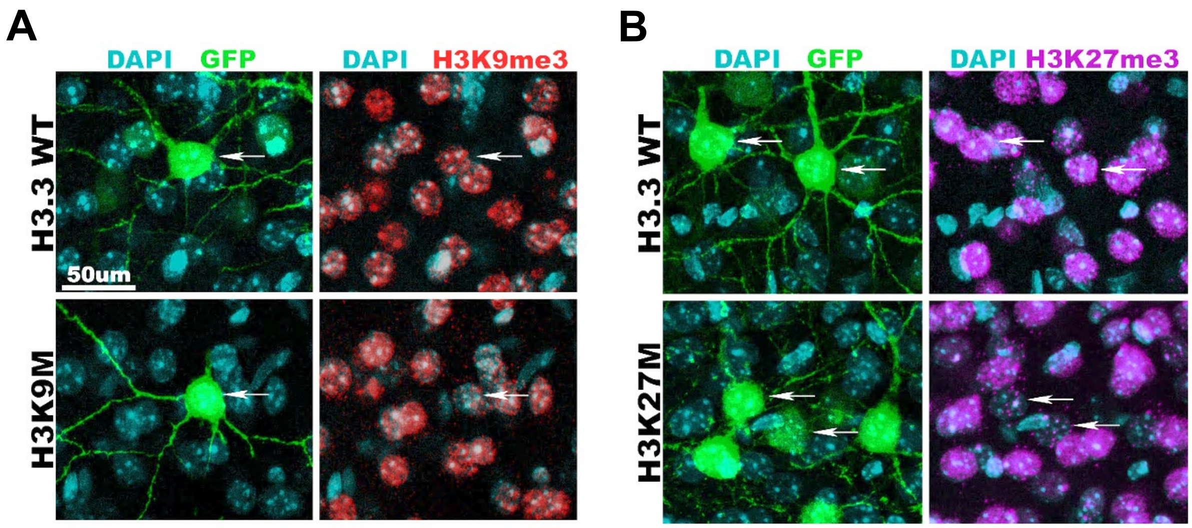


**Fig S6. Inhibition of histone H3 methylation in the adult cortex.** The inducible Cre/FLEX vector system was electroporated at E14.5, followed by tamoxifen administration at P15-P18, and the detection of histone marks at P100. (A) Induction of H3K9M resulted in sustained inhibition of H3K9me3 in comparison to the H3.3 WT-electroporated controls (arrows). (B) The induction of H3K27M led to a decrease of H3K27me3 in $GFP^{+}$ neurons that was maintained at P100 (arrows).
